# An auxin-inducible degron system for conditional mutation in the fungal meningitis pathogen *Cryptococcus neoformans*

**DOI:** 10.1101/2024.11.18.624150

**Authors:** Manning Y. Huang, Matt J. Nalley, Patrick Hecht, Hiten D. Madhani

## Abstract

*Cryptococcus neoformans* is the top-ranked W.H.O. fungal priority pathogen, but tools for generating conditional mutations are limited. Auxin-inducible degron systems permit rapid and effective cellular depletion of a tagged protein of interest upon adding a small molecule. These tools are invaluable, particularly for studying essential genes, which may play important roles in pathogen biology. AID2 is one such system that improves on previous strategies. This system achieves greater sensitivity and specificity using a “bumped” auxin, 5-Ph-IAA, alongside an OsTIR1(F74G) “hole” mutant. We adapted the AID2 system for *C. neoformans* by codon optimizing OsTIR1(F74G) and tested its use in multiple scenarios. We demonstrate that the *C. neoformans* optimized AID2 system enables effective degradation of proteins, including essential proteins, and can be used to discriminate essential from non-essential genes. This tool enables the study of unexplored parts of the *C. neoformans* genome.

## INTRODUCTION

*Cryptococcus neoformans* is an opportunistic fungal pathogen of significant clinical importance (Rajasingham et al. 2017; Fisher and Denning 2023). Our understanding of fungal pathogens has been accelerated by the development of tools for the genetic manipulation of fungal species (Ross and Santiago-Tirado 2024). In *C. neoformans*, a genome-wide deletion collection has permitted the high-throughput dissection of gene function in pathogenesis and basic biological pathways (Boucher et al. 2024 Jan 1). While this collection contains 4328 high-confidence deletion mutants, deletions were not obtained for 2523 genes, representing 36% of the *C. neoformans* proteome. One prominent explanation for an inability to disrupt a given gene of interest is that the gene of interest may be essential. While some proportion of essential genes represent well-conserved pathways, many of these putatively essential genes are uncharacterized with no predicted orthology. As these non-disruptable genes ostensibly drive essential functions in an important human pathogen, additional tools are required for their study.

Controlling gene product expression is an invaluable tool for understanding how essential genes function. Most approaches rely on inhibiting RNA expression to decrease protein expression. This is commonly achieved by replacing the native promoter of a gene of interest with an exogenous promoter to achieve transcriptional repression. Within fungal pathogens, commonly used promoters include the Tet promoter (Tet-On/Tet-Off) and copper repressible promoters (Ory et al. 2004; Park and Morschhäuser 2005). In *C. neoformans*, Beattie and colleagues used the copper repressible promoter *P_CTR4_* to inhibit the essential gene *FKS1*, encoding the catalytic subunit of beta-glucan synthase (Beattie et al. 2022). Inhibition of expression with high copper led to increased cell wall chitin content in response to inhibition. In diploid *C. albicans*, the GRACE strain collection was constructed by systematic deletion of one allele and replacement of the promoter of the second allele with the Tet-Off promoter (Roemer et al. 2003). This collection was used to identify a partial set of *C. albicans* essential genes and identify genes and pathways involved in morphology switching (O’Meara et al. 2015). While effective, promoter replacement strategies present a few disadvantages. Inducible/repressible promoters are known to have “leaky” expression or may overexpress genes under inducing conditions, causing misregulation of essential genes and confusing the interpretation of results. Some essential genes may also have low native RNA expression levels, and repressive promoters might not significantly decrease expression below those levels (Arita et al. 2021). Early methods to influence protein levels without affecting the native promoter include the DaMP system, which generated hypomorphic alleles by disrupting the 3’UTR of a gene of interest, preventing polyadenylation of mRNA and leading to mRNA instability and turnover (Muhlrad and Parker 1999; Schuldiner et al. 2005). The DaMP method was used to construct an essential gene hypomorph library in *S. cerevisiae*, and was used to study biofilm formation in *C. albicans* (Yan et al. 2008; Finkel et al. 2011). Other RNA inhibition-based strategies include RNAi and CRISPRi, which have been adapted to study fungal pathogens (Mouyna et al. 2004; Rappleye et al. 2004; Wensing and Shapiro 2022). However, RNA inhibition indirectly influences protein expression. Furthermore, changes in RNA expression are not strictly correlated with changes in protein expression and ultimately, inhibition efficiency will depend on protein stability and mRNA turnover (Cheng et al. 2018).

Directly targeting a protein of interest for degradation via the ubiquitin-proteasome pathway has recently become increasingly feasible with the development of new methods. These methods include proteolysis targeting chimeras, which use unique fusion proteins to recruit the E3 ligase complex to a tagged protein of interest (PROTACs); dTAGs, which add versatility to PROTACs by using a standardized chimera to target a protein of interest fused with an FKBP12 subunit; and auxin-inducible degrons which behave similarly but use simpler components (Nishimura et al. 2009; Nabet et al. 2018; Prozzillo et al. 2020). In fungal pathogens, auxin-inducible degron systems have been developed for use in *C. albicans* and *C. glabrata* but have yet to be adapted for use in *C. neoformans* (Milholland et al. 2023). Additionally, the development of electroporation-based CRISPR-Cas9 techniques has improved on older biolistics technology for transformation and has made tag-based tools easier to use (Fan and Lin 2018). We previously demonstrated that short homology arms could be used with a *C. neoformans* codon-optimized copy of Cas9, permitting tagging genes of interest with as little as 50 bp of homology (Huang et al. 2022). In this study, we converted and tested the auxin degron AID2 system in *C. neoformans* and demonstrated that this system is robust and tunable to deplete proteins of interest.

## RESULTS

### Design and expression optimization

We set out to create a toolbox for facile inducible protein degradation in *C. neoformans* (Figure 1). We selected the improved AID2 system for this purpose because AID2 is anticipated to cause less leaky degradation and to degrade degron-tagged proteins more completely (Yesbolatova et al. 2020). This system uses an F74G mutant of the Oryza sativa F-box auxin receptor gene *TIR1* (Yesbolatova et al. 2020). When OsTir1 binds auxin, it forms an E3 ligase complex, which ubiquitinates any proteins carrying a specific degron tag, leading to degradation of the protein of interest through the proteasomal degradation pathway (Figure 2A). To implement this system, we synthesized a *C. neoformans* codon-optimized version of *OsTIR1_F74G_* using the codon optimization scheme we previously employed for Cas9 expression (Huang et al. 2022). As introns are required for efficient gene expression in *C. neoformans*, we also included the same efficiently spliced intron we previously used for *CAS9* expression in the optimized *OsTIR1_F74G_* gene (Huang et al. 2022). Previous work in several species has shown that different degron tags can be degraded with varying efficiency (Li et al. 2019; Sepers et al. 2022). Therefore, to maximize degradation efficiency, we chose to test three commonly used degron tags: mAID, mIAA7, and AID*. The mAID and mIAA7 tags, but not the AID* tag, were also optimized for expression using our *C. neoformans* codon optimization scheme.

**Fig 1.**
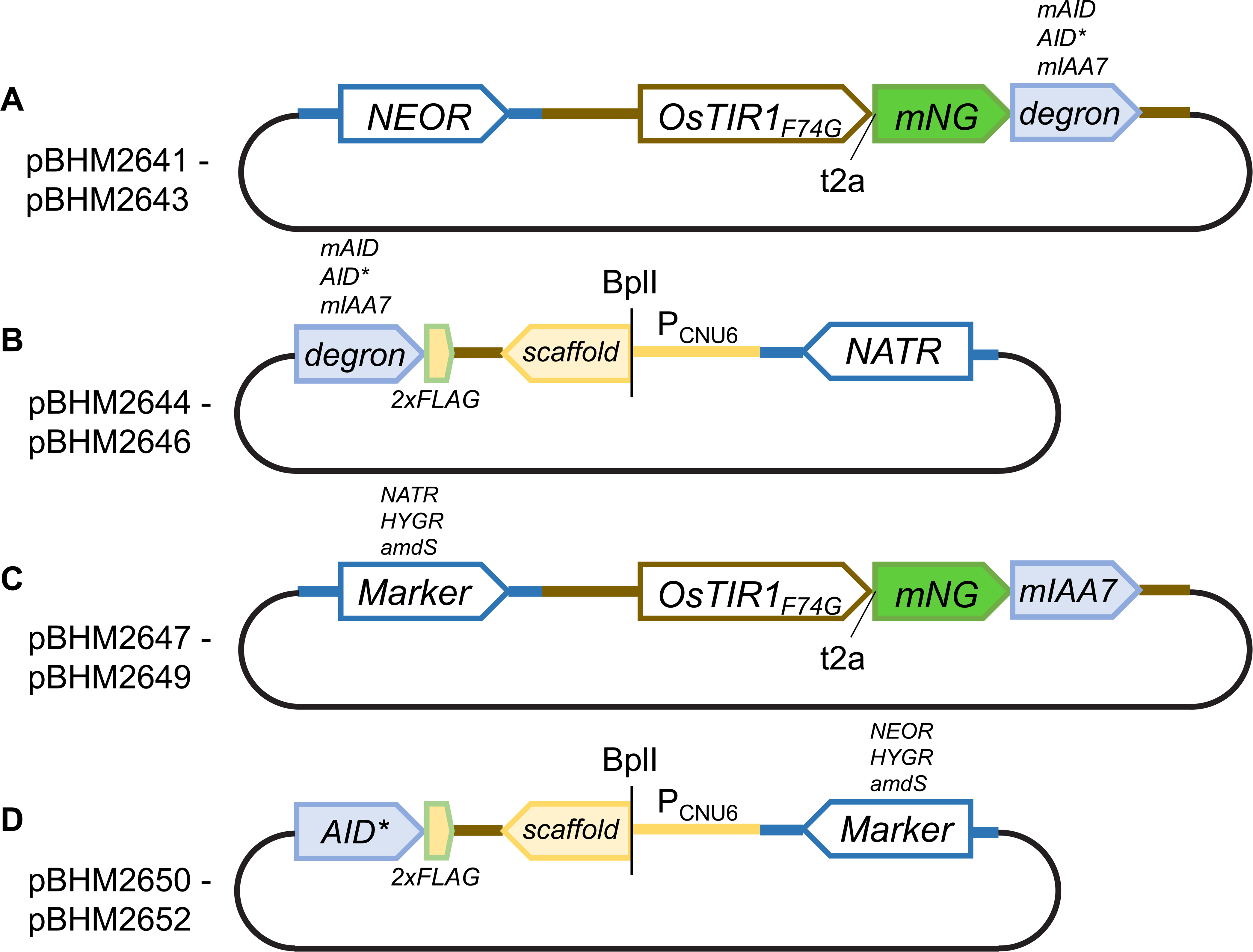
Plasmid Maps. (A) Plasmid series pBHM2641 through BHM2643 carry *OsTIR1-t2a-mNG*- *degron* marked with a neomycin resistance marker for selection and are used to introduce *OsTIR1* at the Safe Haven locus by insertion. Each plasmid carries a different degron, either mAID, AID*, or mIAA7. (B) Plasmid series pBHM2644 through BHM2646 carry only a degron tag (either mAID, AID*, or mIAA7) followed by a 2xFLAG epitope tag, an empty sgRNA, and a nourseothricin resistance marker for transformant selection. These vectors are used as PCR templates to generate PCR products to degron tag a gene of interest. (C) Plasmid series pBHM2647 through BHM2649 are provided for additional selectable marker options and carry *OsTIR1-t2a-mNG*- *mIAA7* but with either the *NATR, HYGR,* or *amdS* markers. (D) Plasmid series pBHM2650 through BHM2652 carry the AID* degron, empty sgRNA, and either *NEOR*, *HYGR*, or *amdS* markers to act as PCR templates for AID* tagging a gene of interest.

**Fig 2.**
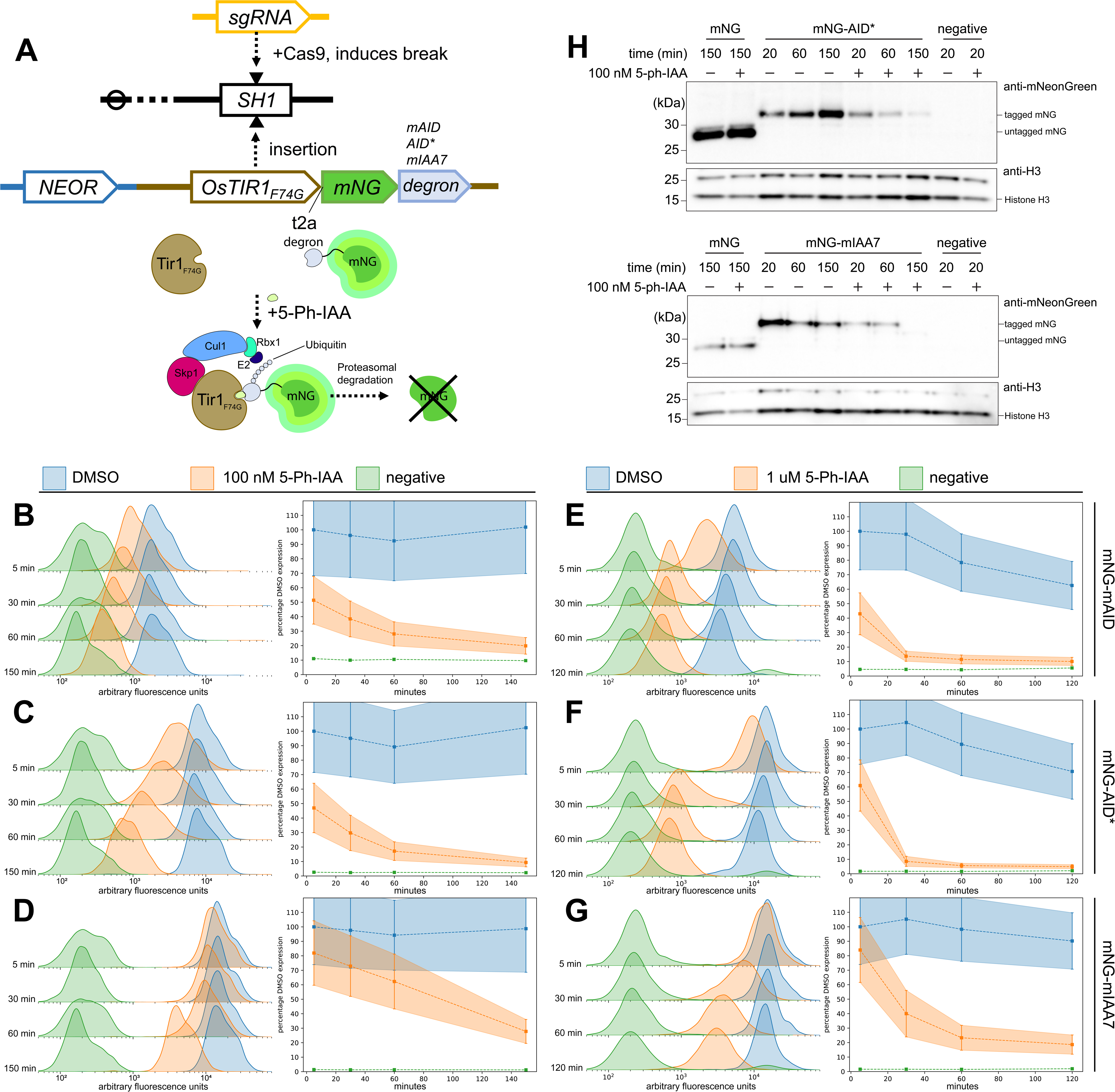
Testing mNeonGreen degradation. (A) Experimental scheme showing insertion of *OsTIR1-t2a-mNG-degron* into the SH1 locus and expected degradation of mNG in the presence of 5-Ph-IAA. (B) Flow cytometry data showing decrease in mNeonGreen signal across multiple timepoints after treatment of an mNG-mAID expressing strain with 100 nM 5-Ph-IAA. Overlapping histograms in left panel show fluorescence signal in arbitrary fluorescence units for different populations of cells. Blue histograms show DMSO treated cells, orange histograms are cells treated with 100 nM 5-Ph-IAA, and green histograms are WT cells not expressing mNG. Each set of 3 histograms correspond to an axis, each axis corresponds to a timepoint – either 5 min, 30 min, 60 min, or 150 min for B-D, or after 120 min for E-G. Right panel summarizes histogram data into a line graph with bars showing IQR with time in minutes on the x axis. Data are presented as a percentage of median DMSO mNG expression at the 5 minute timepoint. (C) Same as B except for mNG-AID*. (D) Same as B except for mNG-mIAA7. (E) Same as B, except cells were treated with 1 uM 5-Ph-IAA. (F) Same as B, except for mNG-AID* treated with 1 uM 5-Ph-IAA. (G) Same as B, except for mNG-mIAA7 treated with 1 uM 5-Ph-IAA. (H) Western blot analysis of mNG signal across multiple timepoints using anti-mNG. Loading controls using anti-H3 are depicted in smaller panels with band corresponding to Histone H3 labeled.

To test if an optimized AID2 system could degrade degron-tagged proteins of interest, we cloned three constructs using the constitutive *TEF1* promoter to drive expression of *OsTIR1_F74G_* followed by a t2a ribosomal skip sequence connected to *mNeonGreen* (*mNG*) tagged with one of the above codon-optimized degrons (Figure 1A). Each construct also carries a G418/neomycin resistance marker for drug selection. We used Cas9 to insert these constructs into the *SH1* locus, and employed flow cytometry to assess fluorescence after treatment of cells with 5-Ph-IAA or DMSO (Arras et al. 2015). In the absence of 5-Ph-IAA, strong fluorescent signal was detected from all three degron-tagged versions of mNG (Figure 2B-G). We observed a relatively reduced signal from DMSO-treated cells carrying mNG tagged with mAID, suggesting that the mAID tag itself might affect mNG expression or destabilize the protein (Figure 2B). However, consistent with expectations for correctly functioning degradation, the addition of 5-Ph-IAA to a final concentration of either 1uM or 100 nM led to rapid loss of mNG signal for cells carrying any of three degron tags (Figure 2B-G). While mAID and AID* tags appeared equally effective at inducing mNG degradation, reducing fluorescent signal to half maximum within 5 minutes, mIAA7 tagged mNG expressing cells showed comparatively slower degradation. As loss of green fluorescent signal could also be explained by quenching of mNG by 5-Ph-IAA, whole cell extracts from AID* and mIAA7 tagged mNG were prepared from cells treated with 100 nM 5-Ph-IAA for western blot analysis using anti-mNG antibodies. We observed reduced band intensity, indicating decreased levels of mNG protein in samples treated with 5-Ph-IAA compared to untreated controls from cells tagged with either mIAA7 or AID* (Figure 2H). These results indicate the *C. neoformans* optimized AID2 system functions robustly and can degrade a tagged protein of interest.

### Auxin-induced depletion of essential proteins

Repeated failure to disrupt a gene of interest is a common approach to identifying essential genes (Akerley et al. 1998; van Opijnen et al. 2009; Segal et al. 2018). However, such an approach is fraught because it draws conclusions from negative data. Auxin degron systems offer an orthogonal approach to test gene essentiality. We selected five genes for further analysis to test the optimized AID2 system in identifying essential genes. These correspond to *HTA1* encoding histone 2A*, CET1*, *TRR1*, *RSA4*, and *ERG8* (standard genome names: *CNAG_06747*, *CNAG_06549*, *CNAG_05847*, *CNAG_04117*, and *CNAG_06001* respectively). Each of these genes except for *HTA1* has been reported as essential in the literature (Missall and Lodge 2005; Ianiri and Idnurm 2015). *CET1*, *TRR1*, *RSA4*, and *ERG8* are essential in *S. cerevisiae* (Giaever et al. 2002). *HTA1* may be nonessential in *S. cerevisiae* because two nearly identical copies exist, while only one copy of *HTA1* exists in the *C. neoformans* genome. *RSA4* essentiality needs to be clarified as the published data disagree, but is likely non-essential as we have been able to disrupt this gene in our efforts to generate a genome-wide deletion collection (Boucher et al. 2024 Jan 1).

We generated plasmids carrying codon-optimized mAID, AID*, and mIAA7 (Figure 1B). Each degron tag is preceded by a GSGSGGSG linker for C-terminal tagging and followed by a *2xFLAG* epitope tag for western blot analysis. This plasmid series also carries an empty sgRNA cassette, with the spacer sequence replaced by a BplI restriction enzyme site. Each plasmid also carries the *NATR* marker for drug selection. While we designed this vector series to be able to use Gibson assembly to clone sgRNAs into each degron plasmid, we used fusion PCR instead to separately amplify sgRNAs targeting downstream regions of our five genes of interest. Plasmids pBHM2644-6 were used as templates for PCR using primers with long overhangs to generate flanking short homology arms on donor DNA carrying the degron tag and *NATR* marker (See Methods). We used CRISPR-Cas9 to generate a *yku80* (encoding the nonhomologous end-joining factor Ku80) insertion mutant derivative from a *TIR1-t2a-mNG-mIAA7* expressing strain to tag all five genes of interest in this strain. Goins and colleagues previously demonstrated that deletion of either *YKU70* or *YKU80* increases rates of homologous recombination in *C. neoformans* (Goins et al. 2006). We therefore decided to use a *yku80* mutant to increase the ease of tagging; we recently demonstrated that transforming a *yku80* disruption with donor and sgRNA cassettes on the same DNA fragment routinely achieves 99%+ efficiency in tagging (Nalley et al., in preparation).

We hypothesized that if a tagged gene was essential, then the tagged strain should fail to grow when degradation was induced by addition of 5-Ph-IAA. In the absence of 5-Ph-IAA, all strains grew similarly to wildtype on YPD media (Figure 3). Independent of the degron version used, *CET1, ERG8, and HTA1* degron-tagged strains all failed to grow on YPD media supplemented with 1 uM 5-Ph-IAA after 24 hours. After 7 days, AID* tagged *CET1* and *H2a* strains continued to show no growth, but mAID and mIAA7 tagged strains grew to varying extent (Figure 3B). For *TRR1*, only AID* and mAID tagged strains showed slightly reduced growth, whereas a mIAA7 tagged *TRR1* strain and showed no growth inhibition. As only one isolate of each degron version was tested per gene of interest, slight differences in growth may not be statistically significant, but our results still demonstrate that depletion efficiency can depend on the degron version used and the tagged gene of interest. Failure to induce sufficient degradation may explain the slightly sick phenotype we observed from AID2 depletion of *TRR1*. While in our experiments we used only 1 uM 5-Ph-IAA, a higher concentration of 5-Ph-IAA might drive more complete protein degradation. Overall, our results demonstrate that we were able to successfully identify gene essentiality by depletion and suggest that AID* may be a reliable tag for use in depleting *C. neoformans* genes.

**Fig 3.**
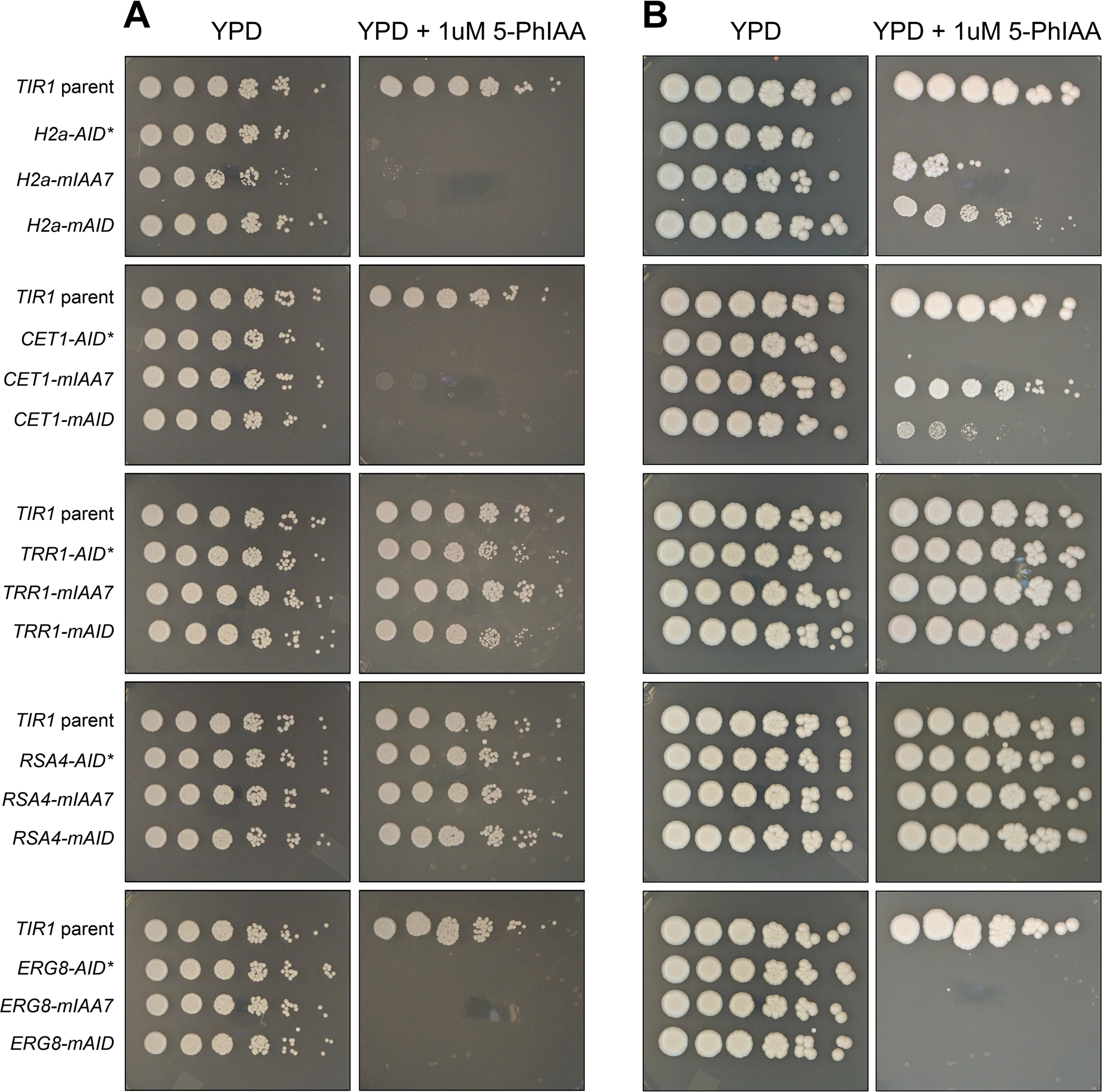
Degron tagged essential genes. Essential genes tagged with either *mAID, AID*,* or *mIAA7* degrons were spotted across a five-fold serial dilution onto YPAD solid media with or without 1 uM 5-Ph-IAA. Cells were grown for either 2 days (A) or 7 days (B) and imaged.

### FKS1 degradation using *C. neoformans* optimized AID2

One potential limitation of auxin degron systems is that membrane-bound or secreted proteins might be inaccessible to *OsTIR1* and may resultantly fail to be degraded. Previous reports have shown success in the depletion of membrane-bound proteins using the AID system in yeast and human cells (Nishimura et al. 2009; Li et al. 2019). These strategies successfully degraded membrane proteins by tagging proteins of interest on accessible cytoplasmic-facing termini (Nishimura et al. 2009). To test whether the optimized AID2 system could successfully degrade membrane-bound proteins in *C. neoformans* using this strategy, we degron tagged the β-glucan synthase catalytic subunit Fks1. Fks1 is a large transmembrane protein with a cytoplasmic facing C-terminus (Hu et al. 2023) and is clinically relevant to the field as the target of echinocandins, one of the three major classes of antifungals. Previous work by Beattie and colleagues has shown using conditional expression controlled by the copper transporter *CTR4* promoter that *C. neoformans* may tolerate 100-fold reduction in RNA expression with only modest growth defect (Beattie et al. 2022). We successfully generated AID* and mAID tagged versions of *FKS1* and assessed growth in liquid YPD media at 30°C across decreasing concentrations of 5-Ph-IAA. Both AID* and mAID tagged versions of *FKS1* showed greatly reduced growth when compared to untagged or DMSO treated cells, confirming that we had successfully depleted *FKS1* using the *C. neoformans* optimized AID2 system (Figure 4). Additionally for both degron tags, growth rate appeared to follow a dose-responsive relationship between 1 nM and 1 uM 5-Ph-IAA. These results highlight the robustness and tunability of auxin-degron systems.

**Fig 4.**
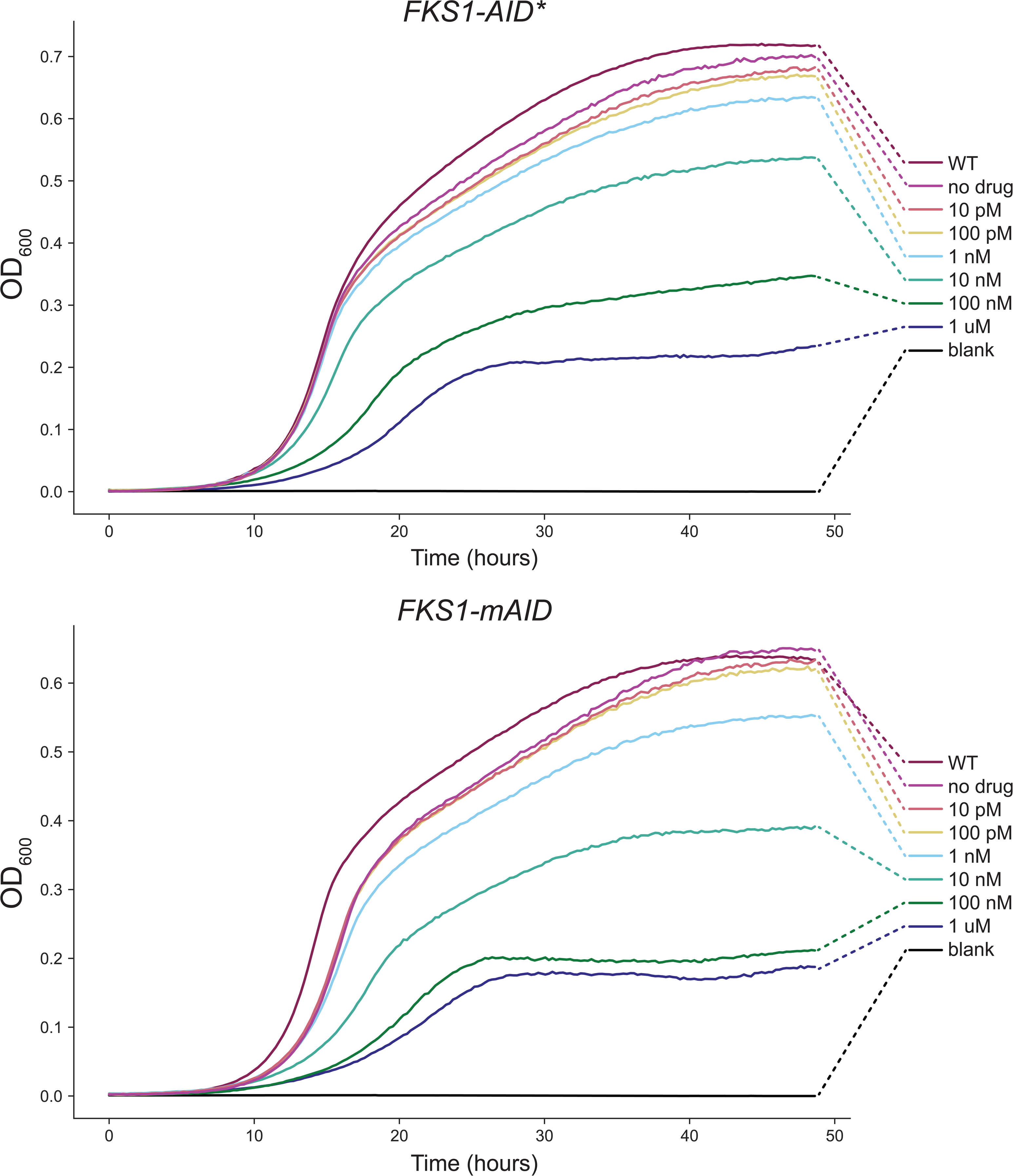
*FKS1* response to depletion. *FKS1-degron* cells corresponding to either *mAID* or *AID\** degron tags were grown in 96 well plates for 48 hours in YPAD supplemented with decreasing concentrations of 5-Ph-IAA. Growth curves show average OD600 measurements from 3 replicates with line coloring indicating the corresponding final concentration of 5-Ph-IAA.

## DISCUSSION

This study presents a valuable addition to the growing *C. neoformans* genetic toolbox. We demonstrate that a *C. neoformans* codon-optimized AID2 degron system is effective for the rapid degradation of proteins of interest. Furthermore, we used the AID2 degron system to tag and deplete essential proteins. Strains with the degron-tagged essential genes *HTA1*, *CET1*, *TRR1*, *ERG8*, and *FKS1* showed severe growth defects when treated with 5-Ph-IAA. We did not observe any phenotypic impact from the depletion of *RSA4*. *RSA4* was identified as an essential gene using segregant analysis and *Agrobacterium-*mediated mutagenesis (Ianiri and Idnurm 2015). However, Billmyre and colleagues identified *RSA4* as non-essential from genome-wide transposon mutagenesis experiments (Billmyre et al. 2024 Jan 1). Additionally, we obtained an *rsa4* deletion mutant in our attempts to generate a *C. neoformans* genome-wide deletion library (Boucher et al. 2024). The disagreement between these results highlights the difficulty of relying on a negative result, namely the failure to delete a gene, to establish gene essentiality. For the same reason, although our results are consistent with *RSA4* being non-essential, degron-based experiments cannot be relied on to establish gene non-essentiality. The simplest explanations would be that we failed to degrade an essential gene sufficiently or that very little expression might be required for viability. For difficult to degrade proteins, a combination of transcriptional and post translational inhibition may provide more reliable control of gene function (Tanaka et al. 2015).

Two critical advantages were described by Yesbolatova and colleagues in the development of the AID2 system - unlike AID, AID2 does not cause leaky degradation, and requires significantly lower concentrations of auxin to induce degradation (Yesbolatova et al. 2020). These advantages were clearly observed in our *C. neoformans* implementation, as degradation was not observed in the absence of 5-Ph-IAA, but could be induced with as little as 1-10 nM 5-Ph-IAA. We observed that the mAID tag caused destablization of mNG in our initial experiments. General wisdom accepts that polypeptide tags may behave idiosyncratically depending on gene and tag utilized. For this reason, we have presented three different tag versions: AID*, mAID, and mIAA7. AID* appears to degrade the most effectively - degrading mNG more quickly than mAID and mIAA7, and also more strongly impacted growth of tagged essential gene strains. We broadly recommend that any *C. neoformans* degron experiments should try at least AID* and one other degron tag version for a gene of interest.

While this work utilized a *yku80* mutant to obtain tagged strains, tag strains can be easily constructed using CRISPR-Cas9 and short homology arms as we have previously demonstrated. However, using a *yku80* strain ensures that nearly all transformants obtained are the correctly tagged strain of interest (Goins et al. 2006) (Nalley et al., in preparation). This approach opens the possibility of generating a genome-wide degron tag collection in either pooled or arrayed formats. We recently used pooled chemical-genetic interaction mapping to functionalize many previously uncharacterized genes, but this experiment was limited by the genes within the *C. neoformans* deletion collection (Boucher et al. 2024 Jan 1). A degron tag collection could include essential genes and genes with severely sick deletion phenotypes, increasing the mutant phenotypes we might observe and thereby improving the resolution of chemical-genetic interaction mapping. We anticipate that methods based on degron techniques will open interesting avenues of study in this important pathogen.

## METHODS

### Media and growth conditions

*C. neoformans* strains were routinely grown in liquid YPD media (2% w/v Bacto Peptone (#211820, Gibco), 2% w/v dextrose, 1% w/v yeast extract (#212720, Gibco)) or on solid YPD media with 2% w/v agar. Where specified, *C. neoformans* strains were also grown in YNB media (0.15% w/v yeast nitrogen base powder without ammonium sulfate (#4027032, MP Biomedicals), 0.5% ammonium sulfate, 2% w/v dextrose). Transformations using amdS selection were plated on YNB without ammonium sulfate plates supplemented with 5 mM acetamide. Liquid cultures were grown at 30°C in a roller drum incubator set to 60 RPM. Cultures grown on solid media were normally incubated at 30°C for 2-3 days unless otherwise specified. To save *C. neoformans* strains, glycerol was added to cells taken from fresh liquid cultures to a final concentration of 15-20% v/v.

5-Ph-IAA (#HY-134653, Med Chem Express) was ordered as solid powder and resuspended to 10 mM in DMSO for stock solution. 5-Ph-IAA stocks and dilutions were kept at - 20°C.

A list of strains, primers, and plasmids used in this study can be found in Table S1.

### Plasmid Cloning

Plasmids were generated by gap repair in *S. cerevisiae*. Briefly, linearized pRS316 vector and DNA products with overlapping homology were transformed into *S. cerevisiae* strain BY4741 using the lithium acetate and ssDNA carrier DNA protocol (Sikorski and Hieter 1989; Gietz and Schiestl 2007). *S. cerevisiae* transformants were plated on YNB plates lacking uracil and plasmid DNA was recovered from yeast transformant cells using the Zymoprep Yeast Plasmid Miniprep II Kit (D2004, Zymo Research), then transformed into DH5α *Escherichia coli* cells by electroporation. Plasmid DNA was recovered from *E. coli* using Nucleospin Plasmid Mini kits (#740588.50, Macherey-Nagel). All plasmids were sequenced by whole-plasmid sequencing performed by Plasmidsaurus using Oxford Nanopore Technology with custom analysis and annotation. All PCRs described in this section were performed using Q5 High-Fidelity Polymerase (#M0491, New England Biolabs).

DNAs containing codon optimized *OsTIR1* and *mAID* were ordered from Genscript in vector format. DNAs containing codon optimized mIAA7 and unoptimized AID* were ordered from IDT as gBlock HiFi Gene Fragments. Two cloning steps were used to assemble each of the plasmids pBHM2461-2463. In the first step, *OsTIR1* was cloned into pRS316 alongside *TEF1* promoter and terminator, as well as a neomycin resistance marker, yielding pBHM2421. This vector also contained additional components which were not ultimately used. Seven PCR products with overlapping homology were used in total and the primers and templates used for each product are listed in Table S1 (Sikorski and Hieter 1989; Arras et al. 2015; Erpf et al. 2019; Huang et al. 2022). In the second round, OsTIR1 and the NEOR marker were amplified from pBHM2421 and cloned into pRS316 with t2a-mNeonGreen amplified from *BLP2:t2a:mNeonGreen C. neoformans* strain CM2034 (Matt Nalley, unpublished data). Four PCR products with overlapping homology were used for this cloning step.

pBHM2426 carries a NATR marker and an empty sgRNA with a BplI cut site between the *C. neoformans U6* promoter and scaffold (Manning Huang, Unpublished Data). Plasmids pBHM2644-6 were cloned from inserting 2 PCR products into pBHM2426 a single step. Primers and products used are listed in Table S1. Note that P23 was paired with one of either P24-26 depending on the degron amplified.

Although the BplI site in pBHM2644-6 was not used for this work, an sgRNA may be cloned into this site. Briefly, 60 bp ssDNA oligos containing a desired sgRNA sequence and flanking homology can be ordered. This oligo may then be directly cloned into BplI digested pBHM2644-6 using an NEB HiFi DNA Assembly Kit (#E2621, New England Biolabs) per manufacturer’s instructions to bridge dsDNA with a ssDNA oligo.

In order to increase the flexibility of using the *C. neoformans* AID2 degron system for tagging, we have additionally generated versions of pBHM2642 with each of the markers *NATR*, *HYGR*, or *amdS* instead of *NEOR*. We have also generated versions of pBHM2644 carrying *HYGR*, *NEOR*, or *amdS* instead of *NATR*.

### Strain Manipulation

Genetic manipulations of *C. neoformans* strains were performed by electroporation as described elsewhere (Lin et al. 2020). Briefly, log phase cells were washed in cold water and treated with 1 mM DTT in electroporation buffer (10 mM Tris-HCl pH 7.5, 1 mM MgCl_2_, 270 mM sucrose) to render cells electrocompetent. DNA was then added to the cells, transferred to a 0.2 cm gap Gene Pulser Electroporation Cuvette (#1652089, Bio Rad) and electroporated in a Gemini X2 Twin Wave Electroporator (#45-2040, BTX Molecular Delivery Systems) with the following settings: 500V, 400Ω, 250 uF. Although we used this specific model, electroporators have easily succeeded with different models but under different optimized settings.

Following electroporation, cells were recovered for 2 hours in liquid YPD media at 30°C with shaking, then plated on solid YPD media supplemented with selective drugs. Final concentrations of either 125 ng/ul nourseothricin (#N51200, RPI) or 200 ng/ul G418 (#61-234-RG, Corning) were used as appropriate.

Primers used to generate sgRNA expressing DNA and donor DNA are listed in Table S1. sgRNA cassettes used in this study were amplified using fusion PCR as previously described using pBHM2329 as template (Huang et al. 2022). All PCRs for this section were run using ExTaq DNA polymerase (#RR001, Takara Bio), cleaned as previously described using Nucleospin Gel and PCR Clean-up kits (#740609, Macherey Nagel) and eluted in water for electroporation into *C. neoformans* cells.

CM2473-5 were generated by transforming CM2049 with linearized pBHM2641-3 respectively without homology arms as well as SH1 sgRNA expressing PCR product to insert *OsTIR1* into the *SH1* locus. CM2476 was generated by transforming CM2474 with *amdS* marker amplified from pPEE7 (Erpf et al. 2019) without homology arms and YKU80 sgRNA expressing PCR product to disrupt *YKU80*. CM2473 served as the parental strain to each essential gene degron tag strain. To generate each degron tagged essential gene strains CM2477-93 used in this study, two PCR products were used, one specifying an sgRNA targeting the downstream region of the gene to be degron tagged and one carrying the degron tag and *NATR* marker. PCRs to amplify degron and NATR marker were run using long oligos with overhangs to provide homology arms. All transformants were verified by colony PCR and PCR genotyping using genomic DNA. Genomic DNA was isolated as previously described using the C. neoformans version of the “smash and grab” protocol (Lin et al. 2020).

### Flow Cytometry

To prepare samples for flow cytometry, *C. neoformans* cells were inoculated in liquid YNB media and grown to OD 0.5. Cells were then split into three culture tubes and either DMSO was added or 5-Ph-IAA was added to a final concentration of 1 uM or 100 nM. Samples were taken from each culture at each time point and were diluted to OD 0.05 with fresh YNB. One ml of the diluted sample was transferred to a round bottom FACS culture tubes and loaded into a BD Accuri C6 Plus. The BD Accuri C6 Plus uses a 488 nm laser with a 533/30 bandpass filter. 10,000 events were acquired per sample on medium speed setting. Events data were analyzed using the python FlowCal library (version 1.3.0) (Castillo-Hair et al. 2016).

### Western Blot Analysis

The same samples used for flow cytometry were also used for Western Blot analysis. Cells were processed as described previously (Boucher et al. 2024 Jan 1). Volume equivalent to two ODs were taken at each time point from mNG-AID* and mNG-mIAA7 samples. Cells were washed in water then resuspended in 10% TCA and incubated on ice. Samples were pelleted at 21000g at 4°C then washed with cold acetone and pelleted again. Pellets were then air-dried and resuspended in a final concentration of 2x NuPAGE LDS sample buffer (#NP0007, Invitrogen), 50mM Tris-HCl pH 8.0, and 100mM DTT along with 0.5 mm zirconia/silica beads (#11079105Z, BioSpec). Samples were bead-beaten using an Omni Bead Rupter Elite with the following settings: 2x90s cycles, 6.0 m/s, 90s rest). Samples were then separated from zirconia beads by centrifugation and saved at -70°C for later western blot analysis.

Before loading, samples were boiled then run on Surepage 4-12% Bis-Tris gels (#M00653, GenScript). Samples were then transferred onto 0.45 um nitrocellulose membranes in Tris-Glycine transfer buffer (25 mM Tris, 250 mM glycine) with 20% methanol. Membranes were blocked for 1 hour in 5% milk in TBST (10 mM Tris-HCl pH 7.4, 150 mM NaCl, 0.1% Tween-20). For primary antibody, membranes were first blotted using 1:1500 ChromoTek mouse-anti-mNeonGreen monoclonal antibody (#32F6, ProteinTech) in blocking buffer overnight at 4°C. Membranes were then washed three times using 15 minute incubations in TBST. Membranes were then incubated for 1 hour at 25°C in 1:8000 goat-anti-mouse IgG (#31430, Invitrogen) HRP-conjugated secondary antibody. Membranes were washed another three times in TBST then visualized using SuperSignal West Pico Plus chemiluminescent substrate (#34580, Thermo Fisher Scientific). Afterwards, membranes were stripped using Restore Western Blot Stripping Buffer (#21059, Thermo Scientific) per manufacturer’s instructions, washed, then steps above were repeated using 1:1000 rabbit-anti-Histone H3 polyclonal antibody (#65-6120, Invitrogen) as primary antibody and 1:8000 goat-anti-rabbit IgG (#65-6120, Invitrogen) as secondary antibody.

### Spotting and Growth Assays

To prepare cells for spotting assays, strains were grown overnight in YPD. For spotting assays, cells were diluted to OD 0.3 in fresh YPD. Five-fold serial dilutions were taken from this culture for a total of 6 dilutions ranging from OD 0.3 to OD 9.6E-5. Three ul of each dilution was spotted onto YPD plates with or without 1 uM 5-Ph-IAA. This roughly corresponds to an expected 9000 down to 3 CFUs per spot from highest to lowest dilution. Cells were grown for two days at 30°C then imaged and allowed to continue to grow at room temperature before being imaged again.

For liquid growth assays, *FKS1* degron tagged strains were first grown overnight in YPD, then serially diluted to OD 0.02 in fresh YPD. 50 ul of YPD with 2x final 5-Ph-IAA concentration to be tested were added to wells of a 96-well untreated round bottom plate (#229590, CELLTREAT). 50 ul of OD 0.02 cells were added to each well except blank control wells. The plate was then covered with a Breathe-Easy polyurethane film (#9123-6100, USA Scientific) following which cells were grown and ODs were measured every 15 minutes across 48 hours in an Tecan Infinite 200 Pro with the following settings: 30°C incubation, orbital shaking with 4 mm amplitude. Four replicates for each degron tagged strain were grown in each 5-Ph-IAA concentration. Growth curve data were analyzed using the Python package Curveball (v0.2.16) (Ram et al. 2019).

## Data Availability

Plasmids pBHM2641-pBHM2652 used in this study were deposited with Addgene. Strains are available upon request.

## Acknowledgements

We thank all the members of the Madhani lab for their insightful discussions that contributed to this work. This work was supported by R01 AI000272.

